# Overcoming Tumor Heterogeneity by Ex Vivo Arming of T Cells Using Multiple Bispecific Antibodies

**DOI:** 10.1101/2021.08.31.458394

**Authors:** Jeong A Park, Nai-Kong V. Cheung

## Abstract

**Purpose:** Tumoral heterogeneity is a hallmark of tumor evolution and cancer progression, being a longstanding challenge to targeted immunotherapy. Ex vivo armed T cells (EATs) using IgG-[L]- scFv bispecific antibodies (BsAbs) are potent tumor-specific cytotoxic effectors. To improve the anti-tumor efficacy of EATs against heterogeneous solid tumors, we explored multi-antigen targeting approaches.

**Methods:** *Ex vivo* expanded T cells were armed with BsAbs built on the IgG-[L]-scFv platform, where an anti-CD3 (huOKT3) scFv was attached to the carboxyl end of both light chains of a tumor specific IgG. Multispecificity was created by combining monospecific EATs, combining BsAbs on the same T cell, or combining specificities on the same antibody. Three multi-antigens targeting EAT strategies were tested: (1) pooled EATs (simultaneous combination of monospecific EATs or alternate EATs (alternating combination of monospecific EATs), (2) dual-EATs or multi- EATs (T cells simultaneously armed with ≥ 2 BsAbs), and (3) TriAb-EATs [T cells armed with BsAb specific for two tumor targets besides CD3 (TriAb)]. The properties and efficiencies of these 3 strategies were evaluated by flow cytometry, *in vitro* cytotoxicity, cytokine release assays, and *in vivo* studies performed in BALB-*Rag2*^-/-^IL-2R-*γc*-KO (BRG) mice xenografted with cancer cell line (CDX) or patient-derived tumor (PDX).

**Results:** Multi-EATs retained target antigen specificity and anti-tumor potency. Cytokine release with multi-EATs in the presence of tumor cells was substantially less than when multiple BsAbs were mixed with unarmed T cells. When tested against CDXs or PDXs, dual- or multi-EATs effectively suppressed tumor growth without clinical toxicities. Most importantly, dual- or multi- EATs were highly efficient in preventing clonal escape while mono- or TriAb- EATs were not as efficient.

**Conclusion:** Arming T cells with multiple BsAbs enabled multi-specific T cell immunotherapy which overcomes tumor heterogeneity without excessive cytokine release.

## INTRODUCTION

T cell immunotherapy has renewed hope for durable cancer cure. However, success has largely been limited to hematological malignancies and a few cancers with high tumor mutational burden. Treatment-related adverse events including cytokine release syndrome, neurotoxicity, and long-term on-target off-tumor toxicities, particularly for targets expressed in normal tissues (e.g. HER2 [1]), are major challenges, hampering their clinical application. For solid tumors, additional hurdles have emerged, such as tumoral heterogeneity, physical barriers, and immunosuppressive tumor microenvironment (TME) [2]. Even for hematologic malignancies highly susceptible to T cell immunotherapy, tumor-associated antigens (TAAs) are often heterogeneous and prone to downregulation or loss, whereby initial responses are not durable and followed by tumor escape and treatment failure [3–5]. To overcome tumor heterogeneity encountered by engineered T cells in solid tumors, increasing specificity to two or more targets has not been adequately explored.

T cell engaging bispecific antibodies (T-BsAbs) have demonstrated promising anti-tumor efficacy in both hematologic malignancies (Blinatumomab, anti-CD19 x anti-CD3)[6] and solid tumors (Catumaxomab, anti-EpCAM x anti-CD3) [7, 8]. Yet despite decades of research and development, only these two BsAbs were clinically approved for cancer treatment. Most T-BsAbs have failed due to insufficient potency or dose-limiting toxicities that were typically cytokine-related. BsAb armed T cells, T cells armed with chemically conjugated anti-GD2 x anti-CD3 [hu3F8 x mouse OKT3 (NCT02173093)], anti-HER2 x anti-CD3 [trastuzumab x mouse OKT3 (NCT00027807)], or anti-EGFR x anti-CD3 [cetuximab x mouse OKT3 (NCT04137536)], have proven safe in multiple clinical trials without cytokine storm, neurotoxicity, or long-term major organ toxicities [9–12]. Recent structure function analyses of BsAbs have shown that ex vivo armed T cells with IgG-[L]-scFv platformed BsAb (EATs) were highly effective against a variety of cancers when compared with those armed with other standard formats of BsAb including chemical conjugates [13–15]. Target antigen-specific EATs effectively infiltrated into tumors despite tissue barriers and immune hostile TME, exerting potent and durable anti-tumor response. To overcome tumor antigen heterogeneity, we now explore multi-antigen targeting approaches. Multispecificity was obtained by combining monospecific EATs, combining multiple BsAbs on the same T cell, or combining specificities on the same antibody, and the following EAT strategies were tested *in vi*tro and *in vivo*: (1) pooled EATs (simultaneous combination of monospecific EATs) or alternate EATs (alternating combination of monospecific EATs), (2) dual- or multi- EATs (T cells simultaneously armed with ≥2 BsAbs), and (3) TriAb-EATs [T cells armed with BsAb specific for two tumor targets besides CD3 (TriAb)].

## METHODS

### 1) Ex vivo T cell activation and arming with BsAb

Peripheral blood mononuclear cells (PBMCs) were separated from buffy coats (New York Blood Center) using Ficoll. The naïve T cells were purified from human PBMC using Pan T cell isolation kit (Miltenyi Biotec, Cat#130096535) and activated and expanded by CD3/CD28 Dynabeads (Gibco^TM^, Cat#11132D) for 7 to 14 days in the presence of 30 IU/mL of IL-2 according to manufacturer’s instructions. T cells were harvested between day 7 and day 14 (median day 10) and, unless stated otherwise, these cultured T cells were used for arming or all T cell experiments. *Ex vivo* armed T cells (EATs) were generated by incubating T cells with T-BsAb for 20 minutes at room temperature. After incubation, these T cells were washed with PBS twice. The T cell number administered per dose was 2x10^7^ cells based on previous reports [40] with supplementary subcutaneous IL-2 (1000 IU).

### 2) Quantification of BsAb bound on EAT

These ex vivo BsAb armed T cells (EATs) were tested for cell surface density of BsAb (MFI) using APC anti-human IgG Fc antibody (Rat IgG2a, κ), (BioLegend, Cat# 410712, RRID:AB_2565790). The MFIs were referenced to antibody binding capacity (ABC) using anti-rat quantum beads Quantum™ Simply Cellular® (QSC) microspheres (Bio-Rad, Cat# FCSC815A, RRID:AB_10061915) for quantification of BsAb bound on T cell.

### 3) Bispecific antibody

All BsAbs were synthesized as previously described (US patent#62/896415) [16–19]. For each BsAb, scFv of huOKT3 was fused to the C-terminus of the light chain of human IgG1 via a C-terminal (G4S)3 linker[20]. N297A and K322A on Fc were generated with site-directed mutagenesis via primer extension in polymerase chain reactions [21]. The nucleotide sequence encoding each BsAb was synthesized by GenScript and subcloned into a mammalian expression vector. Each BsAb was produced using Expi293^TM^ expression system (Thermo Fischer Scientific) separately. Antibodies were purified with protein A affinity column chromatography. The purity of BsAbs was evaluated by size-exclusion high performance liquid chromatography (SE-HPLC) and showed high levels of purity (>90%). The BsAbs remained stable after multiple freeze-thaw cycles. Biochemistry data of the BsAbs used in this study were summarized in the supplementary Table S1 [13, 14].

### 4) Tumor cell lines

Neuroblastoma cell line, IMR-32 (ATCC Cat# CCL-127, RRID:CVCL_0346), osteosarcoma cell line, 143B (ATCC Cat# CRL-8303, RRID:CVCL_2270), primitive neuroectodermal tumor cell line TC-32 (RRID:CVCL-7151), breast cancer cell line HCC1954 (ATCC Cat# CRL-2338, RRID:CVCL_1259), gastric cancer cell line NCI-N87 (ATCC Cat# CRL-2338, RRID:CVCL_1259), acute monocytic leukemia (AML-M5a) cell line MOLM13 (DSMZ Cat# ACC-554, RRID:CVCL_2119), prostate cancer cell line LNCaP-AR (ATCC Cat# CRL-1740, RRID:CVCL_1379), and melanoma cell line M14 (NCI-DTP Cat# M14, RRID:CVCL_1395) were used for experiments. All cancer cell lines were authenticated by short tandem repeats profiling using PowerPlex 1.2 System (Promega, Cat# DC8942), and periodically tested for mycoplasma infection using a commercial kit (Lonza, Cat# LT07-318). The luciferase-labeled melanoma cell line M14Luc, osteosarcoma cell line 143BLuc, and neuroblastoma cell line IMR32Luc were generated by retroviral infection with an SFG-GF Luc vector.

### 5) Antibody Dependent T cell mediated Cytotoxicity (ADTC)

EAT-mediated cytotoxicity was performed using ^51^ Cr release as described previously [18], and EC50 was calculated using SigmaPlot software. Target cell lines were cultured in RPMI-1640 (Cellgro) supplemented with 10% fetal bovine serum (FBS, Life Technologies) and harvested with EDTA/Trypsin. These target cells were labeled with sodium ^51^Cr chromate (Amersham, Arlington Height, IL) at 100 µCi/10^6^ cells at 37^◦^C for 1 hour. After washing twice, these radiolabeled target cells were plated in 96-well plates. EATs were added to target cells at decreasing effector to target cell ratios (E:T ratios), at 2-fold dilutions from 10:1. After incubation at 37^◦^C for 4 hours, the released ^51^Cr was measured by a gamma counter (Packed Instrument, Downers Grove, IL). Percentage of specific lysis was calculated using the formula where cpm represented counts per minute of ^51^Cr released.

<u>100% x (experimental cpm- background cpm)</u> (total cpm-background cpm) Total release of ^51^Cr was assessed by lysis with 10% SDS (Sigma, St Louis, Mo) and background release was measured in the absence of effector cells and antibodies.

### 6) Cytokine release assays

EAT-induced human cytokine release was analyzed *in vitro* and *in vivo*. Human Th1 cell released cytokines were analyzed by LEGENDplexTM Human Th1 Panel (Biolegend, Cat# 741035). Five human T cell cytokines including IL-2, IL-6, IL-10, IFN-γ and TNF-α were analyzed after exposure to target antigen (+) tumor cell lines (*in vitro*). Mouse serum cytokines were analyzed between 3 and 4 hours after each EAT injection.

### 7) *In vivo* experiments

All animal experiments were performed in compliance with Memorial Sloan Kettering Cancer Center’s institutional Animal Care and Use Committee (IACUC) guidelines. *In vivo* anti-tumor response was evaluated using cancer cell line- or patient-derived xenografts (CDXs or PDXs). Cancer cells suspended in Matrigel (Corning Corp, Tewksbury MA) or PDXs were implanted into the right flank of 6–10-week-old BALB-*Rag2*^-/-^IL-2R-*γc*-KO (BRG) mice (Taconic Biosciences) [22]. The following cancer cell lines and cell doses were used: 1x10^6^ of 143BLuc, 5x10^6^ of IMR32Luc, 5x10^6^ of HCC1954, 5x10^6^ of LNCaP-AR, and 5x10^6^ of TC-32. For mixed lineage CDX, 2.5x10^6^ of IMR32Luc and 2.5x10^6^ of HCC1954 were mixed and implanted into each mouse. Three osteosarcoma, one Ewing sarcoma family of tumors (EFT), and one breast cancer PDXs were established from fresh surgical specimens with MSKCC IRB approval. To avoid biological variables, only female mice were used for *in vivo* experiments except LNCaP-AR CDXs using male mice. Treatment was initiated after tumors were established, average tumor volume of 100 mm^3^ when measured using TM900 scanner (Piera, Brussels, BE). Before treatment, mice with small tumors (<50 mm^3^) or infection signs were excluded from randomization to experimental groups. Tumor growth curves and overall survival was analyzed, and the overall survival was defined as the time from start of treatment to when tumor volume reached 2000 mm^3^. To define the well-being of mice, CBC analyses, body weight, general activity, physical appearance, and GVHD scoring were monitored. All animal experiments were repeated twice more with different donor’s T cells to ensure that our results were reliable.

### 8) GD2 by fresh frozen tumor section staining

Fresh frozen tumor sections were made using Tissue-Tek OCT (Miles Laboratories, Inc, Elkhart, IN) with liquid nitrogen and stored at -80°C. The tumor sections were stained with mouse IgG3 mAb 3F8 for GD2 as previously described [23]. Stained slides were captured using a Nikon ECLIPSE Ni-U microscope and analyzed, and the tissue staining intensity and percentage of positive cells were compared with positive and negative controls. Each sample was assessed and graded by 2 independent observers.

### 9) Immunohistochemistry (IHC) for T cell infiltration and HER2 expression

Harvested xenografts were formalin-fixed paraffin-embedded (FFPE) and tested for immunohistochemistry (IHC). IHC staining was performed by Molecular Cytology Core Facility of MSKCC using Discovery XT processor (Ventana Medical Systems). FFPE tumor sections were deparaffinized with EZPrep buffer (Ventana Medical Systems), antigen retrieval was performed with CC1 buffer (Ventana Medical Systems), and sections were blocked for 30 minutes with background buffer solution (Innovex). Anti-CD3 antibody (Agilent, Cat# A0452, RRID: AB_2335677, 1.2μg/mL) and anti-HER2 (Enzo Life Sciences Cat#ALX-810-227-L001, RRID: AB_11180914, 5μg/mL) were applied, and sections were incubated for 5 hours, followed by 60 min incubation with biotinylated goat anti-rabbit IgG (Vector laboratories, cat# PK6101) at 1:200 dilution. The detection was performed with DAB detection kit (Ventana Medical Systems) according to manufacturer’s instruction. All images were captured from tumor sections using Nikon ECLIPSE Ni-U microscope and NIS-Elements 4.0 imaging software.

### 10) Statistics

Statistical analyses of tumor growth and *in vitro* cytokine release were conducted using area under the curves (AUC) to obtain numerical values that integrated all parts of the growth curve of tumors. Two-tailed Student’s t-test was used to determine statistical difference between two sets of data, while one-way ANOVA with Tukey’s post hoc test was used to determine statistical differences among three or more sets of data. All statistical analyses were performed using GraphPad Prism V.8.0 for Windows (GraphPad Software, La Jolla, CA, www.graphpad.com). *P* value < 0.05 was considered statistically significant. Asterisks indicate that the experimental *P* value is significantly different from the controls at * *P* < 0.05; ** *P* < 0.01; *** *P* < 0.001, **** *P* < 0.0001.

## RESULT

### Dual antigens targeting strategies using EAT

We first chose 2 target antigens GD2 (disialogangliosides) and HER2 (human epidermal growth factor receptor 2) to test the efficacy of dual-antigen targeting strategies including pooled EATs (co-administering GD2-EATs and HER2-EATs), alternate EATs (GD2-EATs alternating with HER2-EATs), dual-EATs (T cells simultaneously armed with GD2-BsAb and HER2-BsAb), and TriAb-EATs [T cells armed with trispecific antibody (HER2xGD2xCD3 TriAb)] (Fig.1A).

**Fig. 1.**
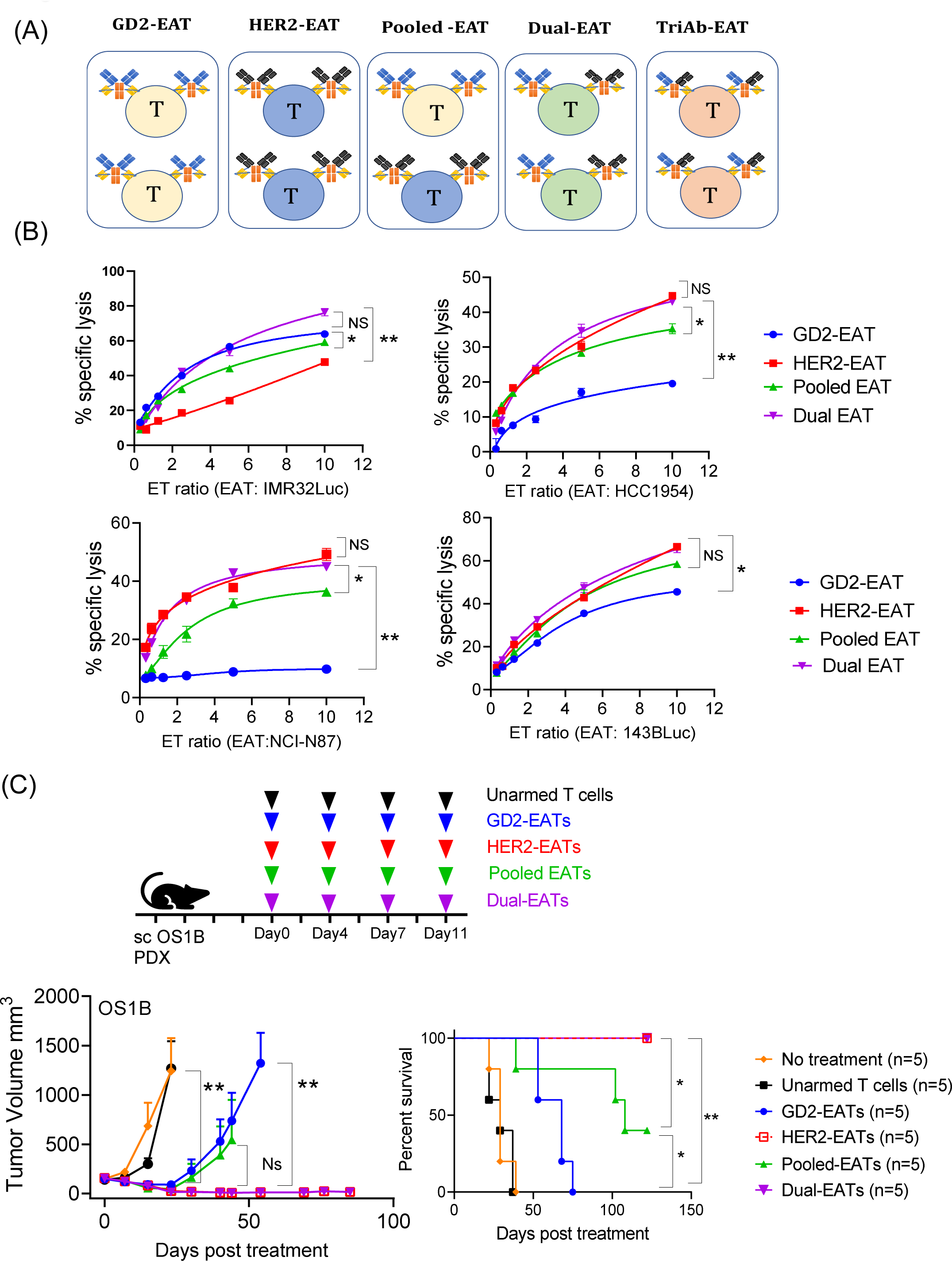
Multi-antigens targeting strategies using *<u>E</u>x viv*o <u>A</u>rmed <u>T</u> cells with IgG-[L]-scFv platformed BsAb (EATs). (A) Representative models of mono-EATs (GD2-EATs or HER2-EATs), pooled-EATs, dual- or multi-EATs, and TriAb-EATs, respectively. (B) *In vitro* cytotoxicity against GD2(+) and/or HER2(+) cancer cell lines was tested and compared among mono-EATs, pooled EATs, and dual- EATs at increasing E:T ratios (effector to target ratio). EATs were armed with 0.5µg of each BsAb per 1x 10^6^ of T cells. GD2(+) IMR32Luc neuroblastoma cell line, HER2(+) HCC1954 breast cancer cell line, HER2(+) NCI-N87 gastric cancer cell line, and both GD2 and HER2 weakly positive (GD2^lo^ HER2^lo^) 143BLuc osteosarcoma cell lines were used respectively. (C) *In vivo* anti- tumor response of mono-EATs [GD2-EATs (10μg of GD2-BsAb/2x10^7^ cells) or HER2-EATs (10μg of HER2-BsAb/2x10^7^cells)], pooled-EATs (5μg/1x10^7^ of GD2-EATs plus 5μg/1x10^7^ of HER2-EATs), and dual-EATs (5μg of GD2-BsAb + 5μg of HER2-BsAb/2x10^7^ cells) was tested against GD2(+) and HER2(+) osteosarcoma PDX (OS1B). Tumor growth curves and overall survival were compared among groups.

First, *in vitro* tumor cell killing by EATs (Fig. 1B) was tested at fixed BsAb arming dose (0.5μg of each BsAb/1x10^6^ T cells) with increasing ET ratios against GD2(+) and/or HER2(+) tumor cell lines (supplementary table S2). Pooled-EATs and dual-EATs showed comparable tumor cell killing when compared with mono-EATs (GD2-EATs or HER2-EATs). While pooled EATs presented an intermediate potency and efficacy between two mono-EATs, dual-EATs showed a similar potency to the target-specific mono-EATs.

*In vivo* anti-tumor effect of multi-EATs was also evaluated using GD2(+) and HER2(+) osteosarcoma PDXs and compared with pooled-EATs (Fig. 1C). While pooled-EATs showed an intermediate anti-tumor effect between two mono-EATs, dual-EATs were equally effective as HER2-EATs; all 5 mice in the dual-EATs or HER2-EATs remained progression-free during follow-up period (up to 150 days post treatment), while none in the GD2-EATs group and only 2 of 5 in the pooled-EATs group showed a long-term remission. We also compared *in vivo* potency of dual-EATs with alternate-EATs using an osteosarcoma 143BLuc CDX mouse model (supplementary Fig. S1). The dual-EATs significantly suppressed tumor growth and showed a comparable anti-tumor effect to HER2-EATs and alternate EATs without increasing toxicities (supplementary Fig. S1).

Next, the anti-tumor efficacy of dual-EATs was compared with TriAb-EATs. We developed a novel GD2xHER2xCD3 trispecific antibody (TriAb) built on the IgG-[L]-scFv platform using a heterodimeric approach as previously described (Fig.2A) [15]. HER2xGD2xCD3 TriAb should engage GD2 and HER2 antigens on tumor cells simultaneously. Their cytotoxicity against multiple cancer cell lines was tested *in vitro* at fixed BsAb arming dose (0.5μg of each BsAb/1x10^6^ T cells) with increasing ET ratios (Fig. 2B). TriAb-EATs were comparable or more effective than GD2- EATs against GD2(+) target cells but were less potent than HER2-EATs against HER2(+) target cells. On the other hand, dual-EATs showed consistently potent cytotoxicity against either GD2(+) and/or HER2(+) cancer cell lines. *In vivo* anti-tumor efficacy of TriAb-EATs was tested using two different osteosarcoma PDX models. Three doses of TriAb-EATs successfully ablated PDX tumors, prolonging survival without obvious toxicity in TEOSC1 PDX model (Fig. 2C). HGSOC1 PDX was more sensitive to GD2-EATs than HER2-EATs, while TriAb-EAT potency was comparable to that of GD2-EATs (supplementary Fig. S2).

**Fig. 2.**
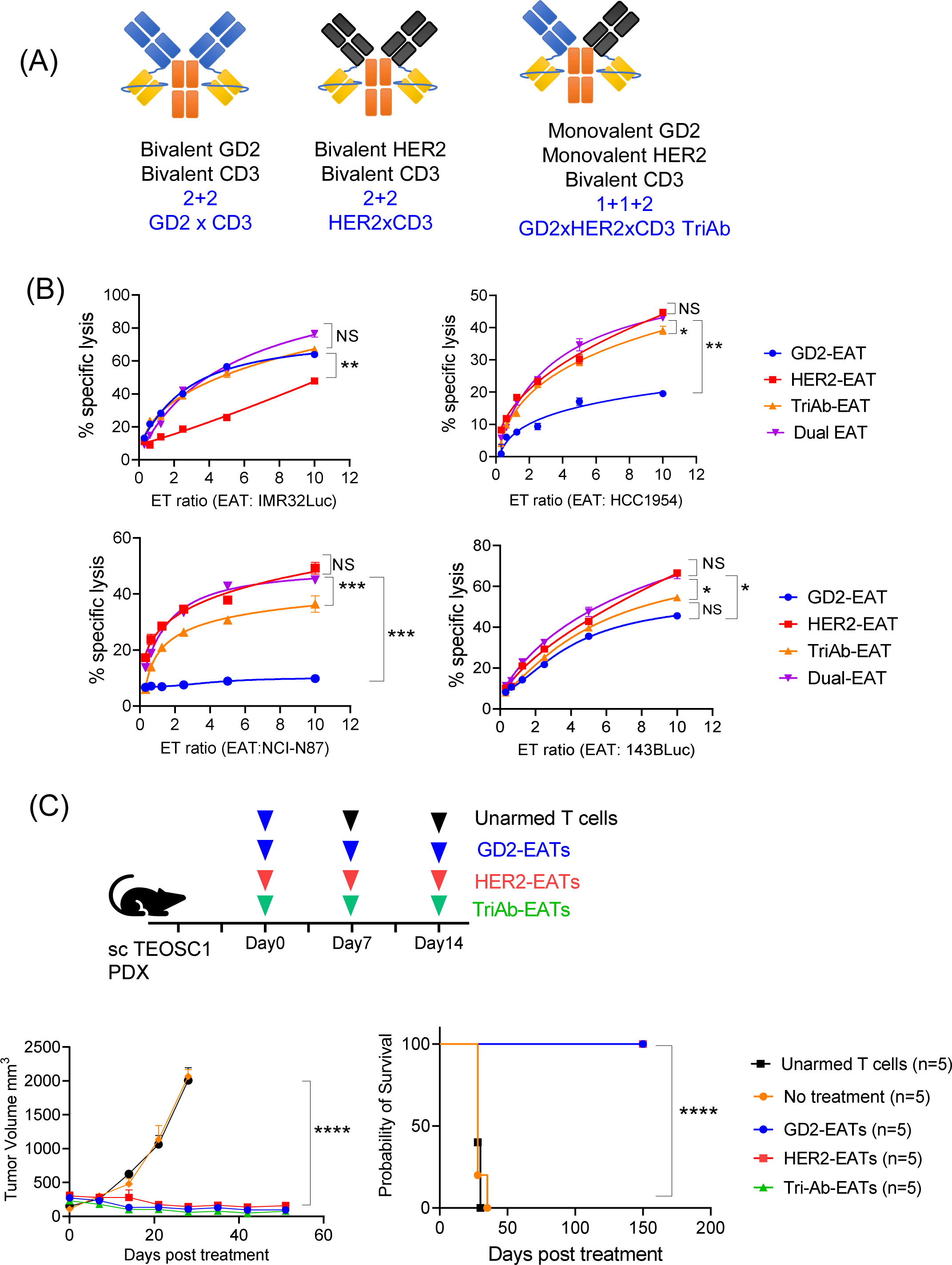
Anti-tumor activity of GD2xHER2xCD3 trispecific antibody (TriAb) armed T cells (TriAb- EATs). (A) Bispecific antibody structure of GD2xHER2xCD3 TriAb. (B) Antibody-dependent T cell- mediated cytotoxicity (ADTC) of TriAb-EAT was compared with mono-EAT (GD2-EAT or HER2-EAT) and dual-EAT against GD2(+) and/or HER2(+) cancer cell lines at increasing E:T ratios. (C) *In vivo* anti-tumor effect of TriAb-EATs against GD2(+) and HER2(+) osteosarcoma PDX (TEOSC1). Three doses of unarmed T cells (2x10^7^ cells) or EATs (10μg of each BsAb/2x10^7^ cells) were administered.

### Optimizing BsAb densities on multi-EATs

Going beyond dual specificities, T cells were simultaneously armed with multiple T- BsAbs specific for GD2, HER2, CD33, STEAP-1, or PSMA, all built on the IgG-[L]-scFv platform. Given the finite CD3 density on human T cells [23], we set out to identify the range and the optimal BsAb surface density as a function of arming dose. Surface BsAb density on EAT was analyzed using anti-human IgG Fc-specific antibody. Precise quantification of BsAb was measured as antibody-binding capacity (ABC) by flow cytometry referenced to quantum beads (Fig. 3A). As the BsAb dose and number have increased, BsAb surface density also has increased. Arming with 5 BsAbs at high arming dose (25µg of each BsAb/10^6^ cells), surface density of BsAb plateaued at approximately 50,000 molecules per T cell.

**Fig. 3.**
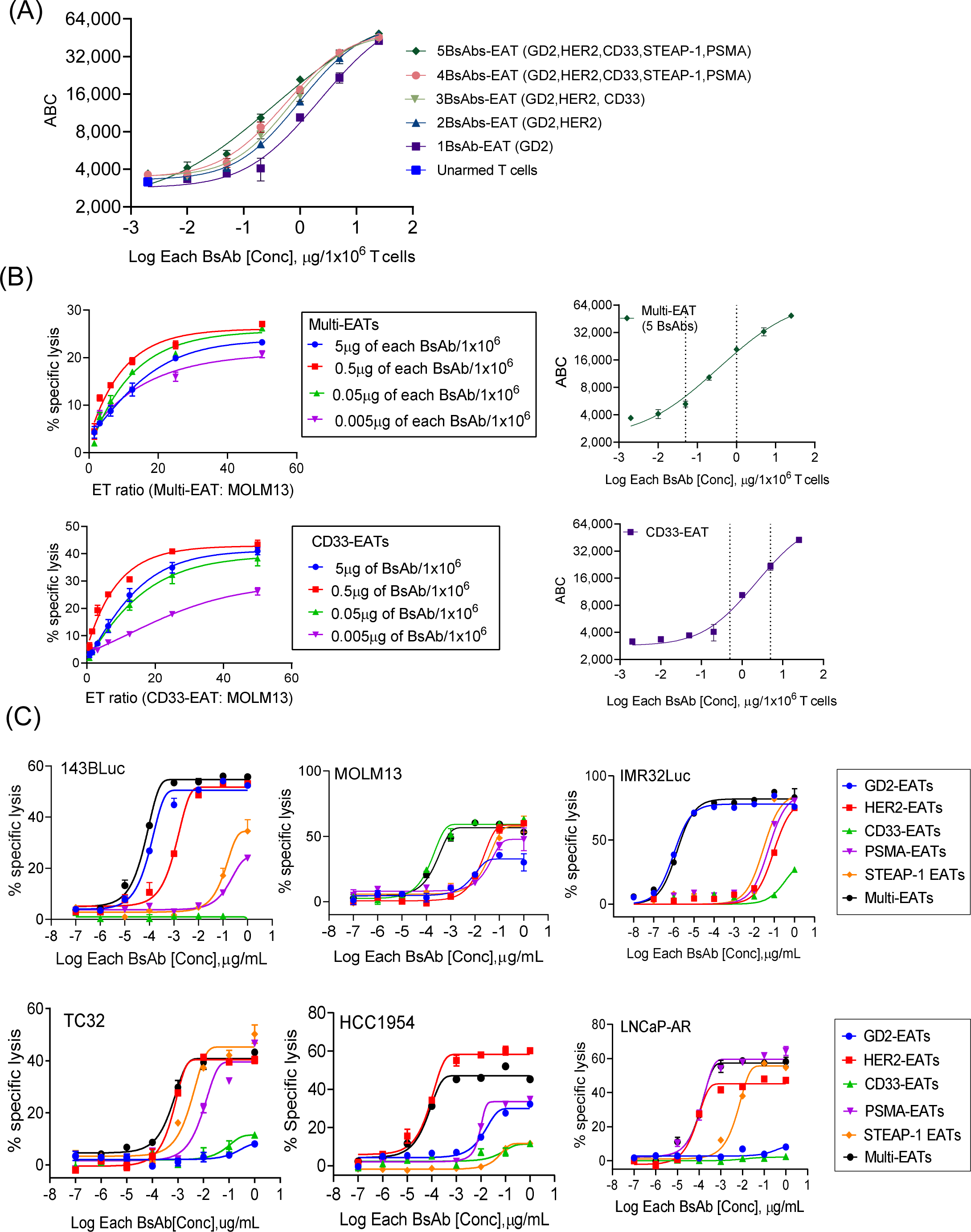
Ex vivo armed T cells with multiple BsAbs (multi-EATs). (a) Surface BsAb density on multi-EAT was analyzed using anti-human IgG Fc-specific antibody and quantified using quantum beads. Geometric mean fluorescence intensities (MFIs) of EATs were measured with increasing arming doses of each BsAb, and BsAb density (MFI) of EAT was referenced to antibody-binding capacity (ABC). (B) *In vitro* cytotoxicity of multi-EATs and CD33-EATs against CD33(+) MOLM13 cell line at increasing E:T ratios and increasing BsAb arming doses. The optimal BsAb densities on T cells were extrapolated from the ADTC assays. (C) *In vitro* cytotoxicity of multi-EATs was tested against a panel of tumor cell lines (E:T ratio was 10:1) and compared with mono-EATs.

To identify the range of optimal surface density of BsAb for multi-EATs, *in vitro* cytotoxicity against CD33(+) leukemia cell line (MOLM13) was studied over a range of ET ratios and BsAb arming doses (Fig. 3B). Multi-EATs (armed with 5 BsAbs each targeting GD2, HER2, CD33, STEAP-1, and PSMA, respectively) showed the best cytotoxicity at the arming dose for each BsAb between 0.05μg/1x10^6^ T cells and 0.5μg/1x10^6^ T cells, corresponding to the BsAb densities between 5,000 and 20,000 molecules per T cell. When referenced to the BsAb density on CD33-EATs which showed the best efficacy between 0.5μg and 5μg of BsAb/1x10^6^ T cells, EATs appear to show the best tumoricidal activity between 5,000 and 20,000 BsAb molecules per T cell.

*In vitro* anti-tumor activity of multi-EATs targeting 5 antigens (GD2, HER2, CD33, PSMA, and STEAP1) was evaluated against varieties of target cell (supplementary table S2) over a range of BsAb arming doses and compared with mono-EATs (Fig. 3C). Multi-EATs exerted consistently the potent anti-tumor activities against each tumor target, comparable to those of target antigen specific mono-EATs, although the maximal cytotoxicity (Emax) did vary depending on the specific targets studied.

#### Ex vivo arming of T cells attenuated cytokine surge from multiple BsAbs

Simultaneous administration of multiple BsAbs could precipitate a cytokine storm. Cytokine release was compared between multi-EATs and multiple BsAbs plus T cells at increasing doses of BsAb (Fig. 4A). Multi-EATs or multiple BsAbs plus T cells were incubated with target cells at 37℃ for 4 hours. Cytokine release of T cells by multiple BsAbs increased by BsAb dose but reached plateaus at 1µg of each BsAb/1x10^6^ cells. The cytokine levels of multi-EATs were significantly lower than those of multiple-BsAbs plus T cells over a range of BsAb doses. When we compared the levels of cytokines released by mono-EATs (HER2-EATs), dual-EATs (HER2/GD2-EATs), triple-EATs (HER2/GD2/CD33-EATs), quadruple-EATs (HER2/GD2/CD33/PSMA-EATs), and quintuple-EATs (HER2/GD2/CD33/PSMA/STEAP1- EATs), the differences were not significant among groups (Fig. 4B). Although IL-2, IL-10, IFN-γ, and TNF-α levels increased with BsAb arming dose, there was no excessive cytokine release with additional BsAbs for multi-EATs.

**Fig. 4.**
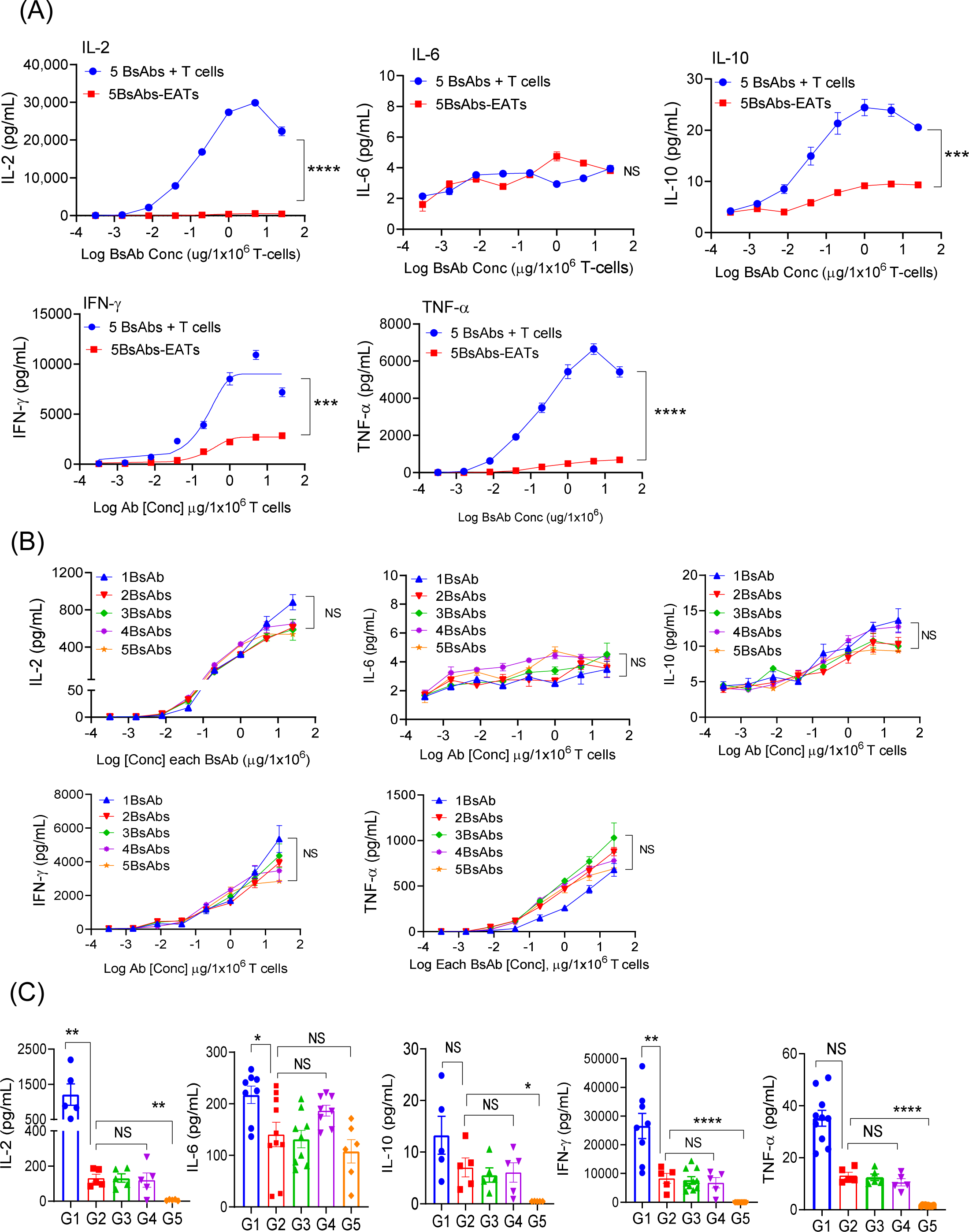
Cytokine release by multiple BsAbs plus T cells and multi-EATs. (A) TH1 cell cytokines (IL-2, IL-6, IL-10, IFN-γ, and TNF-α) were measured in the supernatants after 4 hours of incubation of 5 BsAbs plus T cells or 5 BsAb armed T cells (5BsAbs-EATs) with target cells at increasing doses of each BsAb (0.0003µg/1x10^6^ cells to 25µg/1x10^6^ cells). Mixture of multiple cancer cell lines consisting GD2(+) M14Luc, HER2(+) HCC1954, CD33(+) HL60, PSMA(+) LNCaP-AR, and STEAP1(+) TC32 were used as target cells. ET ratio (effector to target cell ratio) was 20:1. (B) *In vitro* cytokine release of multi-EATs was compared following an increase in the number of BsAb. ET ratio was 20:1, and mixture of multiple cancer cell lines consisting GD2(+) M14Luc, HER2(+) HCC1954, CD33(+) HL60, PSMA(+) LNCaP-AR, and STEAP1(+) TC32 were used as target cells. (C) *In vivo* TH1 cytokine levels were analyzed 4 hours after second dose of EAT in GD2^lo^HER2^lo^ 143BLuc osteosarcoma cell line xenograft (CDX) mouse model. G1, GD2-BsAb and unarmed T cells; G2, multi-EATs (GD2/HER2/CD33/PSMA/STEAP1-EATs); G3, GD2-EATs; G4, HER2-EATs; G5, unarmed T cells. BsAb dose and T cell number were fixed at 10µg for each BsAb and 2x10^7^ for T cell per injection.

*In vivo* cytokine levels by multi-EATs were also analyzed post treatment and compared among groups (Fig. 4C). Multi-EATs (50μg of total BsAb/2x10^7^ cells, G2) released significantly less IL-2, IL-6, IFN-γ, and TNF-α than directly injected GD2-BsAb (10μg) plus unarmed T cells (2x10^7^ cells) (G1), and there was no significant difference in cytokine release among GD2-EATs (G3), HER2-EATs (G4), and multi-EATs (G2).

### Multi-EATs as multi-specific cytotoxic T lymphocytes

#### In vivo anti-tumor properties against diverse cancer types

*In vivo* anti-tumor effect of multi-EATs was tested against diverse cancer xenografts representing different tumor types (Fig. 5A). Multi-EATs (2μg of each BsAb x 5 BsAbs/2x10^7^ T cells per injection) significantly suppressed tumor growth and consistently showed competitive anti-tumor effect to mono-EATs against a panel of target appropriate cancer xenografts, including HER2(+) M37 breast cancer PDX, PSMA(+) LNCaP-AR prostate cancer CDX, GD2(+) IMR32Luc neuroblastoma CDX, and STEAP1(+) ES3a EFT PDXs (Fig. 5B), without clinical toxicities. For IMR32Luc CDXs, multi-EATs exerted a robust anti-tumor effect surpassing the efficacy of GD2-EATs, significantly prolonging survival.

**Fig. 5.**
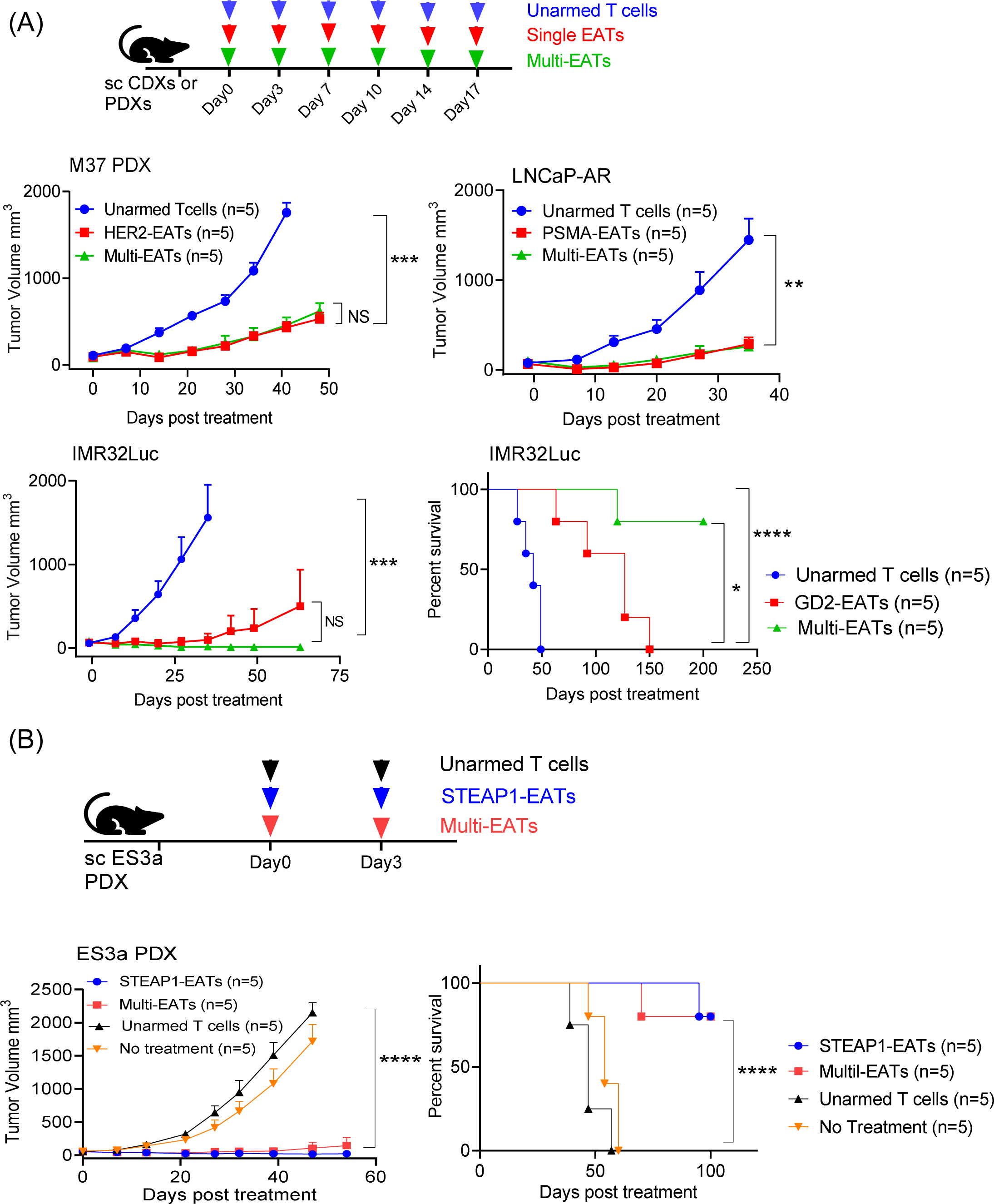
In vivo anti-tumor activities of multi-EATs. (A) In vivo anti-tumor effect of multi-EATs was tested against a variety of cancer xenografts including M37 breast cancer PDXs, LNCaP-AR prostate cancer CDXs, and IMR32Luc neuroblastoma CDXs. Six does of EATs or unarmed T cells were administered. (B) In vivo anti- tumor effect of multi-EATs was compared with single antigen targeted STEAP1-EATs against Ewing sarcoma family of tumor (EFT) PDXs. Two doses of EATs or unarmed T cells were administered. BsAb dose and T cell number were fixed at 2µg for each BsAb and 2x10^7^ for T cell per injection.

#### Multi-EATs were highly effective against tumor models with antigen heterogeneity

We next studied the ability of multi-EATs to overcome tumor heterogeneity by creating a mixed lineage, i.e., GD2(+) IMR32Luc mixed with HER2(+) HCC1954 (1:1 ratio) (Fig. 6A).

**Fig. 6.**
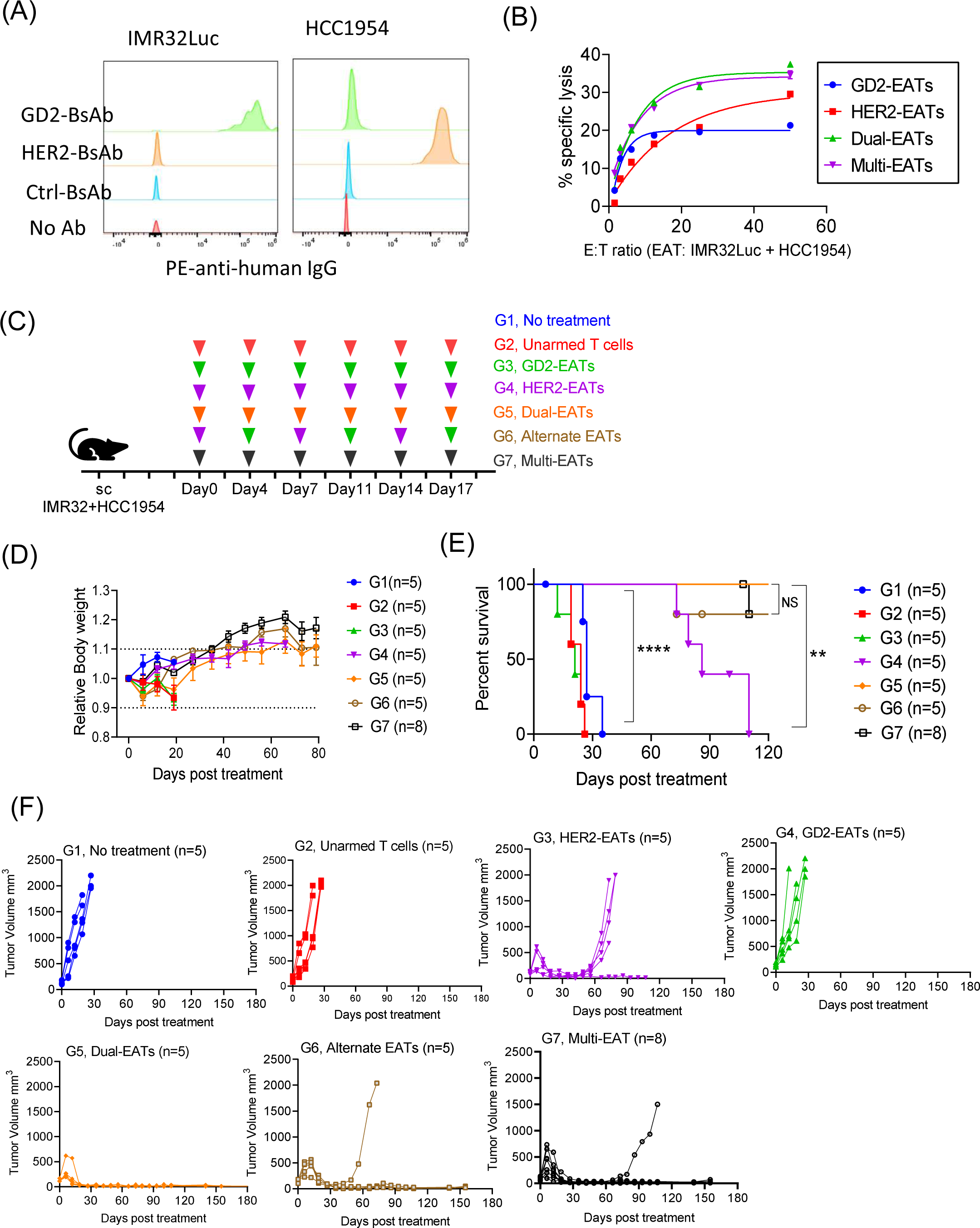
Anti-tumor efficacy of multi-EATs against mixed lineage targets. (A) Antibody binding intensities (MFIs) of each cancer cell line. (B) *In vitro* cytotoxicity was tested against IMR32Luc and HCC1954 mixed lineage. (C) Schematic overview of treatment for IMR32Luc and HCC1954 mixed lineage xenografts using multi-antigen targeting EAT strategies. BsAb dose and T cell number were fixed at 10µg for each BsAb and 2x10^7^ for T cell per injection. (D) Mouse body weight during follow-up period. (E) Overall survival by treatment. (F) Tumor response by treatment groups.

Dual-EATs (T cells armed with GD2-BsAb and HER2-BsAb) and multi-EATs (T cells armed with 5 BsAbs targeting GD2, HER2, CD33, PSMA, and STEAP1, respectively) induced stronger cytotoxicity against this mixed lineage than GD2-EATs or HER2-EATs (Fig. 6B). This enhanced *in vitro* cytotoxicity of dual- or multi-EATs translated into more potent *in vivo* anti-tumor response. A mixture of two cell lines was xenografted subcutaneously and treated with dual- or multi-EATs and compared efficacy with mono-EATs (Fig. 6C). EATs were fixed at 10µg of each BsAb/ 2x10^7^ T cells for each injection. No clinical toxicities were observed and there was no weight loss throughout the follow-up period (Fig. 6D). While GD2-EATs or HER2-EATs failed to produce durable responses against this mixed lineage CDX, dual-, multi-, or alternate-EATs successfully regressed tumors, producing long-term survival (Fig. 6E and 6F). Dual- or multi- EATs both surpassed the efficacy of mono-EATs, significantly improving tumor-free survival (vs. HER2-EATs, *P*=0.0033; vs. GD2-EATs, *P*<0.0001). There was no significant difference in efficacy among dual-, multi-, and alternate- EATs.

We also tested the efficacy of TriAb-EATs against this mixed lineage. While TriAb-EATs showed enhanced *in vitro* cytotoxicity compared with GD2-EATs or HER2-EATs, it was not as effective when compared to dual- or multi-EATs (supplementary Fig. S3). *In viv*o anti-tumor activity of TriAb-EATs was also tested against this mixed lineage CDXs (supplementary Fig. S3). Tumors regressed following TriAb-EATs, but the response was not durable: all 5 mice relapsed after short-term response, contrasting to dual- or multi-EATs where long-term disease-free survival extended past 140 days in 3 out of 5 and 4 out of 5 mice, respectively.

#### Multi-EATs overcame tumoral heterogeneity: Histologic response of mixed lineage CDX to multi-EATs

The mixed lineage CDXs were harvested after treatment and analyzed their antigen expression. Gross examination of these tumors presented distinct differences between IMR32Luc and HCC1954 lineages (Fig. 7A and suplementary Fig. S4A). Following treatment with GD2- EATs or TriAb-EATs tumors grossly resembled HCC1954 CDXs, while following HER2-EATs tumors appeared IMR32Luc CDXs. After treatment with alternate-EATs, dual-EATs, or multi- EATs tumors had appearances of a cross between IMR32Luc and HCC1954 CDXs, while untreated tumors or those treated with unarmed T cells more resembled HCC1954 CDXs, consistent with rapid outgrowth of HCC1954 overtaking IMR32Luc. H&E staining results corresponded to their gross phenotypes (Fig. 7B and suplementary Fig, S4B). While following treatment with GD2-EATs or TriAb-EATs histology revealed poorly-differentiated invasive ductal breast carcinoma, following treatment with HER2-EATs tumors showed immature, undifferentiated, small round neuroblasts accompanied by Homer-Wright pseudo-rosettes, typical characteristics of neuroblastoma. With no treatment or treatment with unarmed T cells, or with recurrence after initial response to alternate-, dual- or multi-EATs, tumors presented mixed lineage with a slight prominence of breast cancer features. Fresh frozen tumor staining with anti-GD2 antibody (hu3F8) (Fig. 7C and supplementary Fig. S4C) and FFPE tumor staining with anti-human HER2 antibody (Fig. 7D and supplementary Fig. S7D) also showed contrasting results following treatment. While the tumors without treatment or treated with unarmed T cells showed heterogeneous staining patterns, those treated with GD2-EATs or TriAb-EATs became GD2 negative and HER2 strongly positive; vice versa, those tumors treated with HER2-EATs were strongly GD2 positive and HER2 negative. Mono-EATs successfully ablated target antigen positive tumor cells but did not affect target antigen negative clones. On the other hand, the escaped tumors following treatment with dual-, alternate-, or multi-EATs were both GD2 and HER2 weakly positive. Total target antigen loss seen with mono-EATs was not observed in tumors treated with these multi-antigen targeting EATs, allowing repeat response to multi-EATs (supplementary Fig. S5).

**Fig. 7.** Analysis of tumor response by immunohistochemical (IHC) staining. (A) Gross phenotypes of tumors in each treatment group: a, unarmed T cells; b, GD2-EATs; c, HER2-EATs; d, TriAb-EATs; e, alternate EATs; f, dual-EATs; g, multi-EATs. (B) H&E staining of tumors in each treatment group: a, unarmed T cells; b, GD2-EATs; c, HER2-EATs; d, TriAb- EATs; e, alternate EATs; f, dual-EATs; g, multi-EATs. (C) Fresh frozen tumor staining with anti-human GD2 antibody (hu3F8): a, unarmed T cells; b, GD2-EATs; c, HER2-EATs; d, TriAb- EATs; e, alternate EATs; f, dual-EATs, g, multi-EATs. (D) IHC staining of formalin-fixed paraffin-embedded (FFPE) tumor sections with anti-human HER2 antibody: a, unarmed T cells; b, GD2-EATs; c, HER2-EATs; d, TriAb-EATs; e, alternate EATs; f, dual-EATs; g, multi-EATs. (E) IHC staining of FFPE tumor sections with anti-human CD3 antibody: a, unarmed T cells, b, GD2-EATs; c, HER2-EATs; d, TriAb-EATs; e, alternate EATs, f, dual-EATs; g, multi-EAT.

## DISCUSSION

To test the hypothesis that T cells endowed with multiple specificities could overcome tumor heterogeneity preventing treatment resistance or clonal escape, we explored the potential of multi-antigen targeting EAT strategies. T cells simultaneously armed with multiple BsAbs built on the IgG-[L]-scFv platform retained tumor selectivity and anti-tumor potency both *in vitro* and *in vivo* without excessive cytokine release. Multi-EATs targeting GD2, HER2, CD33, PSMA, and STEAP1 demonstrated identical and in some cases robust anti-tumor efficacy compared with mono-EATs against individual tumor targets. More importantly, while mono-EATs could ablate tumors in an exquisitely antigen-specific manner and were unable to control antigen negative clones in the mixed lineage tumor system, dual- or multi-EATs could overcome tumor heterogeneity and target antigen loss, two major limitations of single antigen targeted T cell immunotherapy. Multi-EATs were more effective than pooling of monospecific EATs or multiplying specificities on the same protein molecule, with the added advantages of being a plug- N-play system offering simplicity in manufacture.

Most of human cancers show heterogeneous antigen expression, and single antigen targeted approaches are rarely curative. Furthermore, the TAAs undergo downregulation, mutation, or loss under selective immune pressure following T cell immunotherapy [24]. Multi-antigen targeted strategies have the potential to overcome these antigen escape mechanisms. Simultaneous targeting CD19/CD20 or CD19/CD20/CD22 using “OR” logic-gated tandem CAR T cells [25] reduced or prevented target antigen escape, offering an advantage over single CAR T cells or pooled populations of monospecific CAR T cells [26–29]. Given the minimal requirement of BsAb (only 500 to 5,000 molecules) per T cell for anti-tumor activity [13], multiple BsAbs built on the same IgG-[L]-scFv platform can be installed on each T cell before the maximum capacity is reached (30,000 to 56,000 molecules per T cells [13, 23]). Since T cell loading is mediated through the same anti-CD3 scFv in IgG-[L]-scFv constructs, BsAb surface density should be predictable and consistent. By adjusting the arming doses, the relative density of each BsAb on each T cell could be fine-tuned and optimized. The same Boolean logic for “OR” and “AND” gates for CAR T cell [25, 30] can be applied to multi-EATs, using appropriately designed BsAbs that can either activate or inhibit T cell function. In our studies, dual- or multi-EATs showed a synergistic anti-tumor effect when simultaneously encountering multiple antigens. The formation of bi- or multi- valent immune synapses when dual- or multi-EATs exposed to heterogeneous tumors co-expressing multiple TAAs would be crucial to exert synergistic anti-tumor effect and prevent antigen escape [27].

Besides tumor heterogeneity, antigen loss or down-regulation has been another challenge to immunotherapy [3]. CD19 loss or mutation following CD19-CAR T cells, or CD22 density dwindling after CD22-CAR T cell therapy are associated with treatment resistance or relapse [31–34]. To address this issue, dual antigen targeting strategies, such as CD19/CD22 dual-specific CAR T cells, pooling of CD19- and CD22- CAR T cells, or sequential treatment with CD19- and CD22- CAR T cells, have been explored with variable success [35]. Our data support an alternative approach using proteins instead of genes to expand T cell specificity. In contrast to the tumors treated with mono-specific GD2-EATs or HER2-EATs forcing target antigen loss, the relapsed tumors following alternate-, dual- or multi-EATs therapy retained their target antigen expression, and escaped EFT PDXs after multi-EAT therapy responded to rechallenges, implicating a major advantage over conventional single antigen targeted immunotherapy.

One of the concerns of multi-antigen targeted T cell immunotherapies is on-target off-tumor toxicities. On-target off-tumor toxicities following the infusion of CAR T cells or BsAbs can cause serious or life-threatening adverse effects, and the extent and severity of toxicity could be amplified by increasing numbers of targeting antigens. Although we did not observe any additional toxicity related to multi-EATs, the mouse models we used have major limitations because of the specificity of BsAbs for human not mouse antigens. High target antigen affinity could increased the severity of on-target off-tumor toxicities, while reducing target antigen affinity to a certain threshold (Kd < 10^−8^ M) significantly decreased toxicities without affecting anti-tumor efficiency of T cells [36]. CAR T cells with µM affinity retained strong antitumor activity while lowering off-tumor toxicities than their nM affinity counterparts [37], suggesting that avidity optimization is an effective strategy to reduce on-target off-tumor recognition by multivalent targeted immunotherapy [38, 39]. Multi-EATs take advantage of avidity-affinity balance with the potential to reduce the on-target off-tumor side effects while expanding the spectrum of responsive tumor types. But more importantly, while multi-specific CAR T cells are life-long and such toxicities could be prolonged and life-threatening, EATs have limited functional life expectancy; as the BsAbs get metabolized, T cells should revert to their nonspecific states, alleviating the risk of life-threatening long-term toxicities. Akin to multi-agent chemotherapy, multi-EATs have the potential to increase potency, reduce toxicity, and overcome tumor heterogeneity [4, 5]. This area of research deserves further exploration and optimization in order to confront cancer resistance.

## Acknowledgement

We would like to thank Dr. Hong Xu for providing the anti-HER2 BsAb and performing binding kinetic studies for BsAbs, Dr. Sayed Shahabuddin Hoseini for anti-CD33 BsAb, Dr. Steven Tsung-Yi Lin for the anti-STEAP1 and anti-PSMA BsAb, Dr. Brian H. Santich for creating the GD2xHER2xCD3 trispecific antibody. We also want to acknowledge Hong-fen Guo for flow cytometry analysis of tumor cell lines and her expertise in the preparation and biochemical analyses of BsAbs, Yi Feng for staining fresh frozen tumor tissues with hu3F8 antibody, and Drs. Sarah Katz and Ileana Miranda in the Laboratory of Comparative Pathology for reviewing pathology of the xenografts. We thank Dr. Elisa De Stanchina for providing PDXs for these studies. This work was supported by funds to NKC from Enid A. Haupt Endowed Chair, the Robert Steel Foundation, and Kids Walk for Kids with Cancer. Technical service provided by the MSK Animal Imaging Core Facility, Antitumor Assessment Core Facility, and Molecular Cytology Core Facility were supported in part by the NCI Cancer Center Support Grant P30 CA008748.

## Author contributions

Jeong A Park and Nai-Kong V. Cheung designed the experiments, interpreted and analyzed the results and wrote the manuscript. All authors reviewed this manuscript and approved final the manuscript.

## Disclosure of Conflicts of Interest

Both NKC and JP were named as inventors on the patent of EATs filed by MSK. Both MSK and NKC have financial interest in Y-mAbs, Abpro-Labs and Eureka Therapeutics. NKC reports receiving commercial research grants from Y-mabs Therapeutics and Abpro-Labs Inc. NKC was named as inventor on multiple patents filed by MSK, including those licensed to Ymabs Therapeutics, Biotec Pharmacon, and Abpro-labs. NKC is a SAB member for Eureka Therapeutics.

## Abbreviations

ABC: Antibody binding capacity of the cell Alternate
EAT: Alternately administered EAT
AUC: Area under the curve
BRG: BALB-Rag2-/-IL-2R-γc-KO
CAR: Chimeric antigen receptor
CDX: Cell line-derived xenograft
Dual-EAT: T cells simultaneously armed with 2 BsAbs
EAT: Ex vivo armed T cells with IgG-[L]-scFv platformed BsAb
ET: ratio Effector to target cell ratio
FFPE: Formalin-Fixed Paraffin-Embedded
GD2: Disialogangliosides
GVHD: graft-versus-host disease
HER2: Human epidermal growth factor receptor 2
IACUC: Institutional animal care and use committee
IHC: Immunohistochemistry
Mono-EAT: Single antigen targeted EAT
Multi-EAT: T cells simultaneously armed with multiple BsAbs
PBMC: Peripheral blood mononuclear cells
PDX: Patient-derived xenograft
Pooled-EATs: Combination of more than 2 EATs each armed with target specific BsAb
PSMA: Prostate-specific membrane antigen
SE-HPLC: Size-exclusion high performance liquid chromatography
STEAP1: Six transmembrane epithelial antigen of the prostate 1
T-BsAb: Tumor microenvironment
TriAb: BsAb specific for two tumor targets besides CD3
TriAb-EAT: T cells armed with trispecific antibody

---

**Supplementary Table S1.**
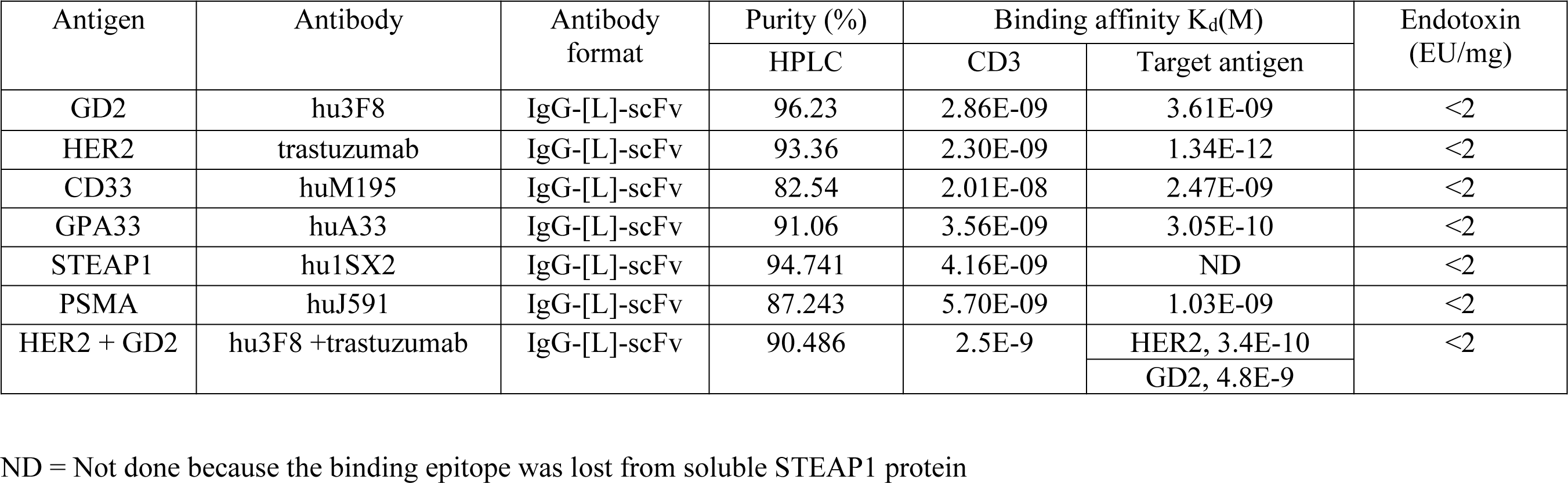
Purity, affinity and endotoxin of each bispecific antibody preparations.

**Supplementary Table S2.** Bispecific antibody binding (MFI) to tumor cell lines by flow cytometry (MFI).

| Cancer | Cell line | Unstained control<br>(MFI) | GD2<br>(MFI) | HER2<br>(MFI) | CD33<br>(MFI) | PSMA<br>(MFI) | STEAP1<br>(MFI) |
| --- | --- | --- | --- | --- | --- | --- | --- |
| Neuroblastoma | IMR32Luc | 80 | 3870 | 78 | 78 | 76 | 78 |
| Osteosarcoma | 143BLuc | 92 | 656 | 1131 | 129 | 138 | 139 |
| Melanoma | M14Luc | 78 | 4685 | 235 | 80 | 79 | 76 |
| Gastric cancer | NCI-N87 | 110 | 101 | 15019 | 102 | 101 | 101 |
| Breast cancer | HCC1954 | 105 | 199 | 5187 | 92 | 1027 | 1719 |
| Ewing sarcoma family of tumors | TC32 | 87 | 147 | 339 | 76 | 71 | 301 |
| Prostate cancer | LNCaP-AR | 103 | 182 | 800 | 138 | 1027 | 1719 |
| Acute myeloid leukemia | MOLM13 | 124 | 209 | 138 | 3546 | 137 | 362 |
|  | HL60 | 103 | 116 | 113 | 5981 | 110 | 111 |

**Supplementary Fig. S1.**
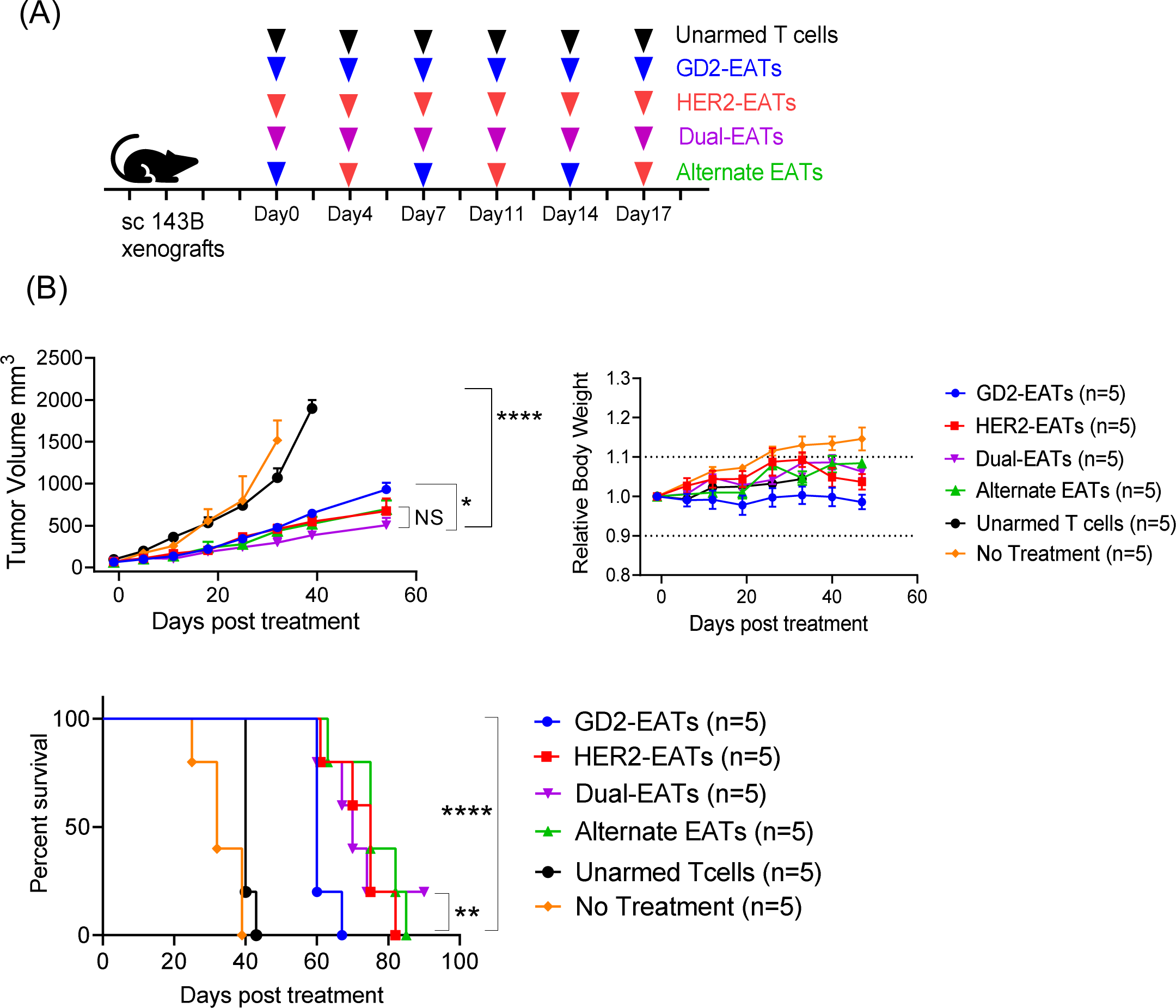
In vivo anti-tumor efficacy of dual- or alternate EATs. (A) Schematic overview of treatment. Six doses of unarmed T cells or EATs were administered intravenously into GD2^lo^HER2^lo^ osteosarcoma 143BLuc cell line xenograft (CDX). BsAb dose and T cell number were fixed at 10µg for each BsAb and 2x10^7^ for T cell per injection. Alternate EATs were given by administering GD2-EATs and HER2-EATs alternately. (B) In vivo anti-tumor response was compared among groups. Tumor growth, body weight of mice during follow-up period, and overall survival were plotted and compared among groups.

**Supplementary Fig. S2.**
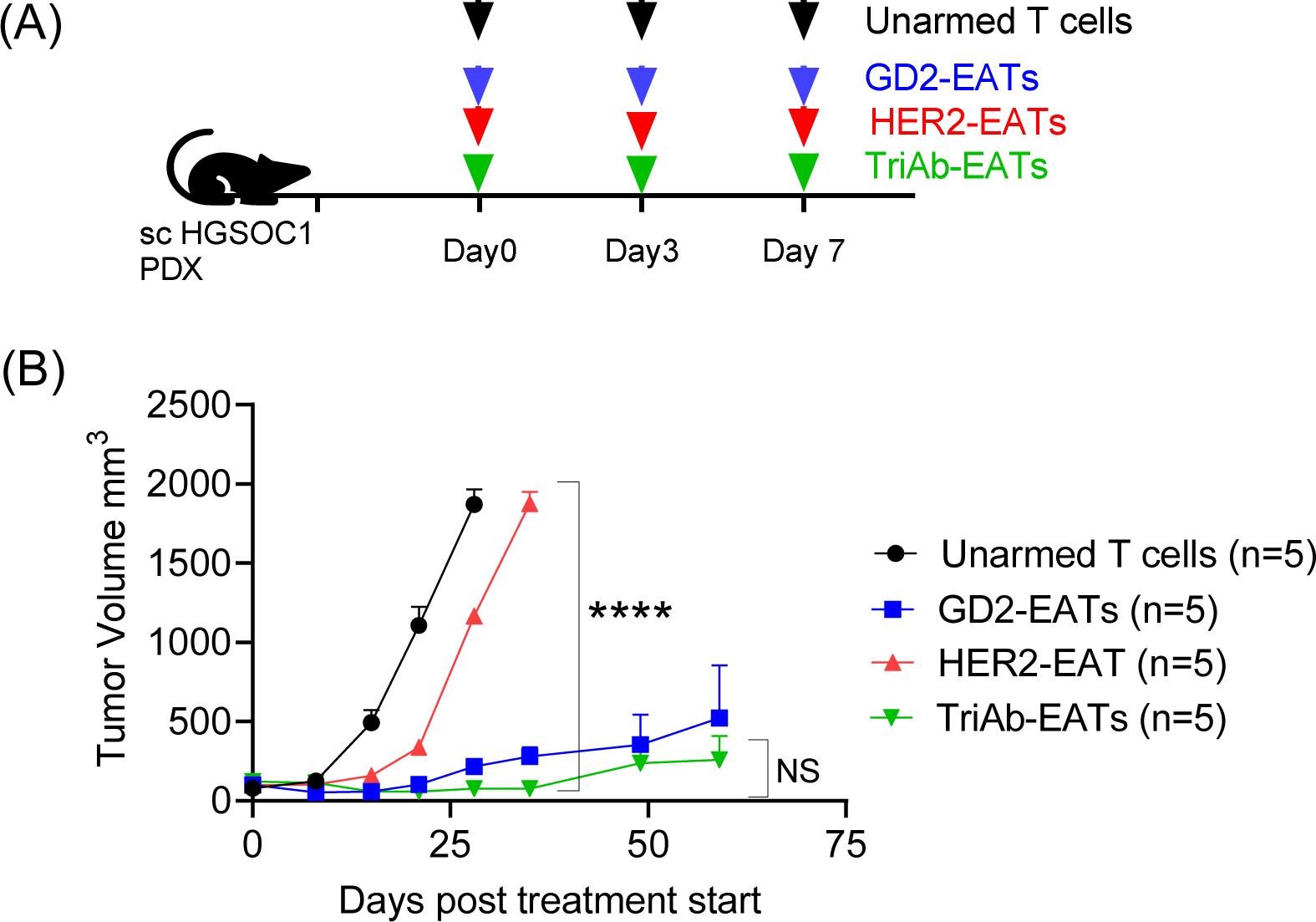
*In vivo* anti-tumor effect of TriAb-EATs. (A) Schematic overview of treatment. Three doses of unarmed T cells or EATs were given intravenously into osteosarcoma PDXs (HGSOC1). 2x10^7^ of unarmed T cells or EATs (10µg of each BsAb/ 2x10^7^ of T cell) were administered iv twice per week. (B) *In vivo* anti-tumor response was compared among groups.

**Supplementary Fig. S3.**
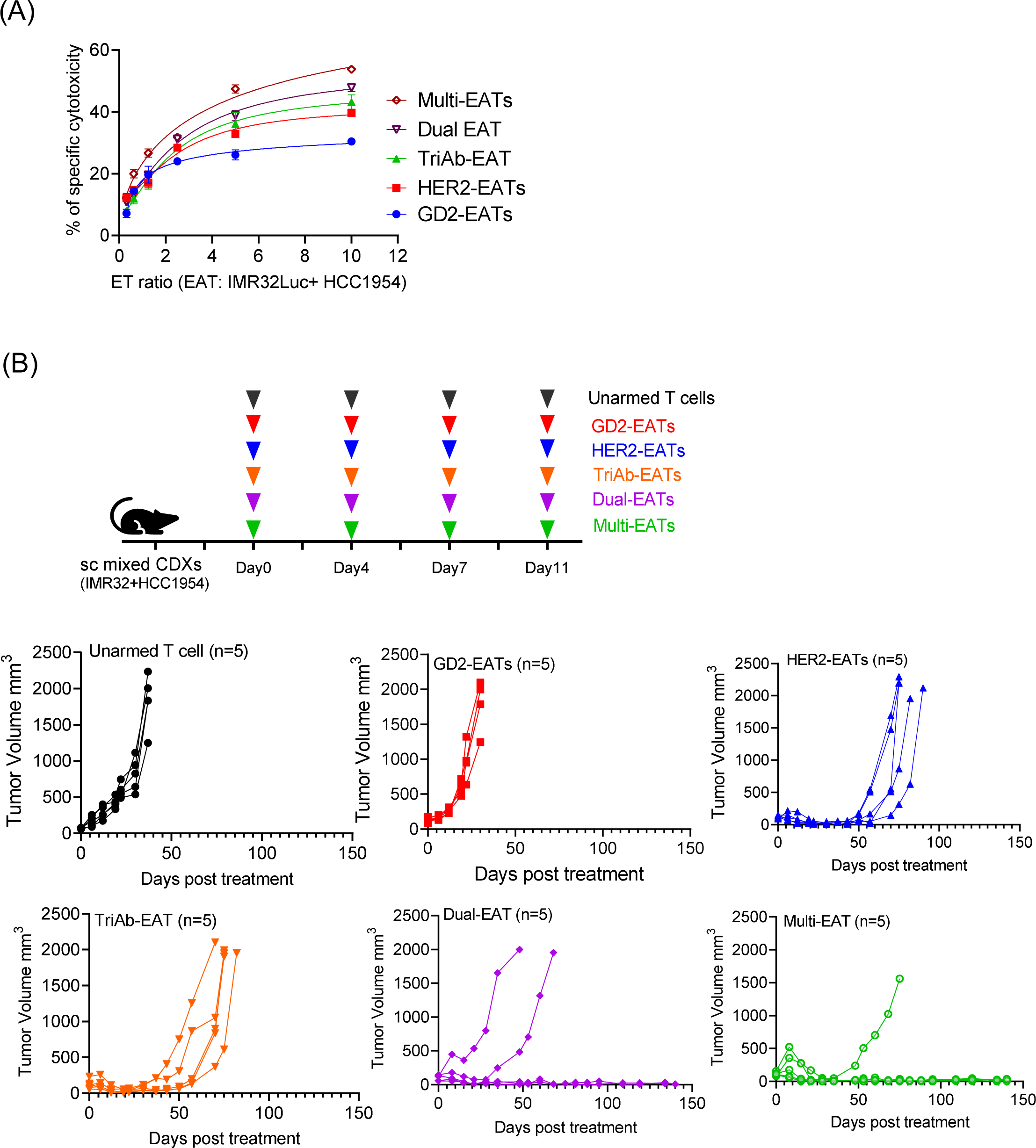
*In vitro* and *in vivo* anti-tumor activity of multi-EATs against mixed lineage. (A) *In vitro* cytotoxicity of multi-EATs was tested against GD2(+)IMR32Luc and HER2(+) HCC1954 mixed lineage and compared with TriAb-EATs and mono-EATs. (B) Schematic overview of treatment for IMR32Luc and HCC1954 mixed lineage xenograft using multiple EAT strategies. (D) *In vivo* anti-tumor activity of multi-EATs was compared among groups including TriAb-EATs. BsAb dose and T cell number were fixed at 10µg for each BsAb and 2x10^7^ for T cell per injection.

**Supplementary Fig. S4.**
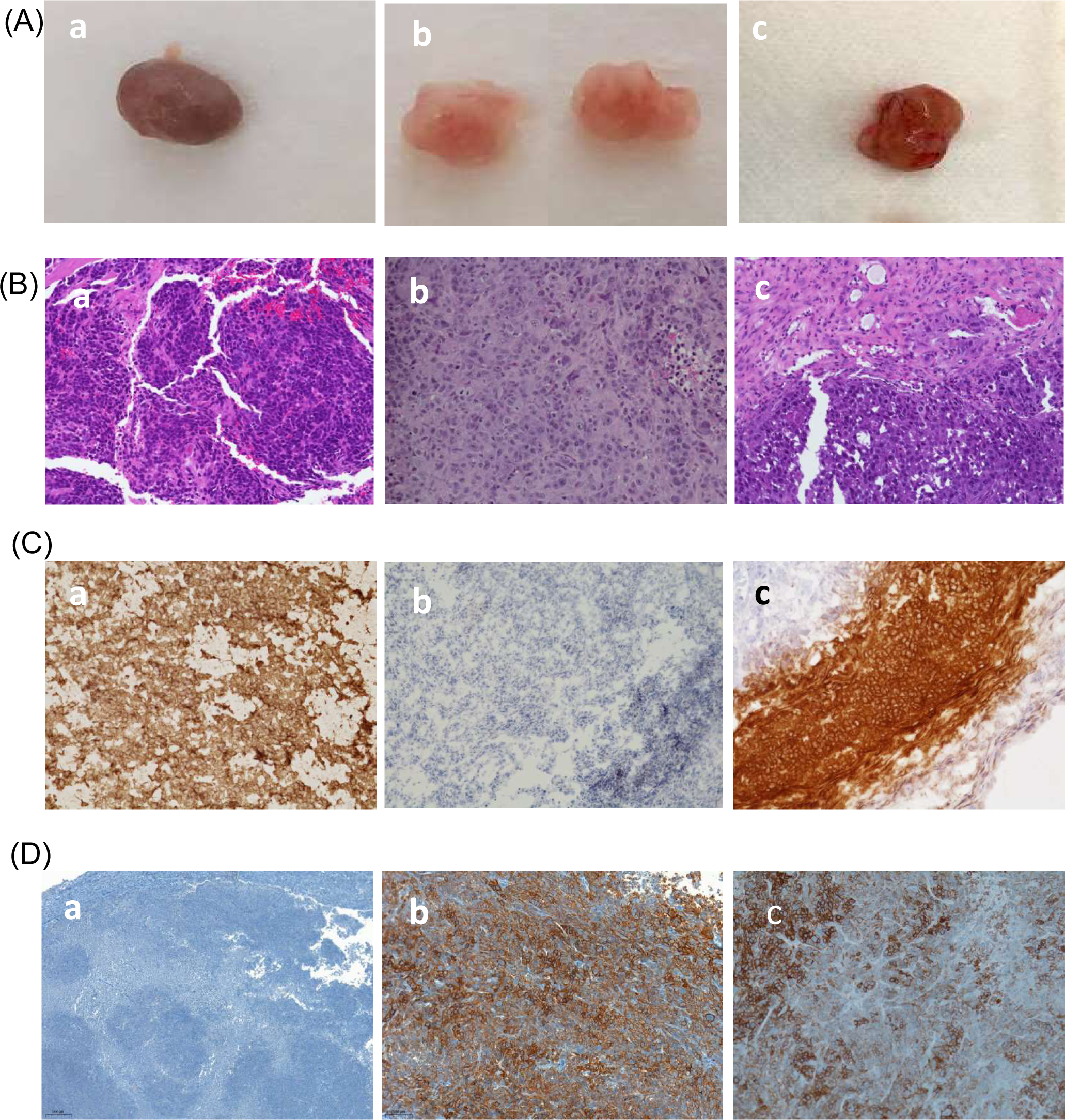
Histologic features of IMR32Luc-, HCC1954-, and IMR32Luc and HCC1954 mixed lineage- xenografts. (A) Gross phenotypes of tumors: a, IMR32Luc cell line xenograft (CDX); b, HCC1954 CDX; c, IMR32Luc and HCC1954 mixed lineage CDX. (B) H&E staining of tumors: a, IMR32Luc CDX; b, HCC1954 CDX; c, IMR32Luc and HCC1954 mixed lineage CDX. (C) Fresh frozen tumor staining with anti-human GD2 antibody (hu3F8): a, IMR32Luc CDX; b, HCC1954 CDX; c, IMR32Luc and HCC1954 mixed lineage CDX. (D) IHC staining of formalin-fixed paraffin-embedded (FFPE) tumor sections with anti-human HER2 antibody: IMR32Luc CDX; b, HCC1954 CDX; c, IMR32Luc and HCC1954 mixed lineage CDX.

**Supplementary Fig. S5.**
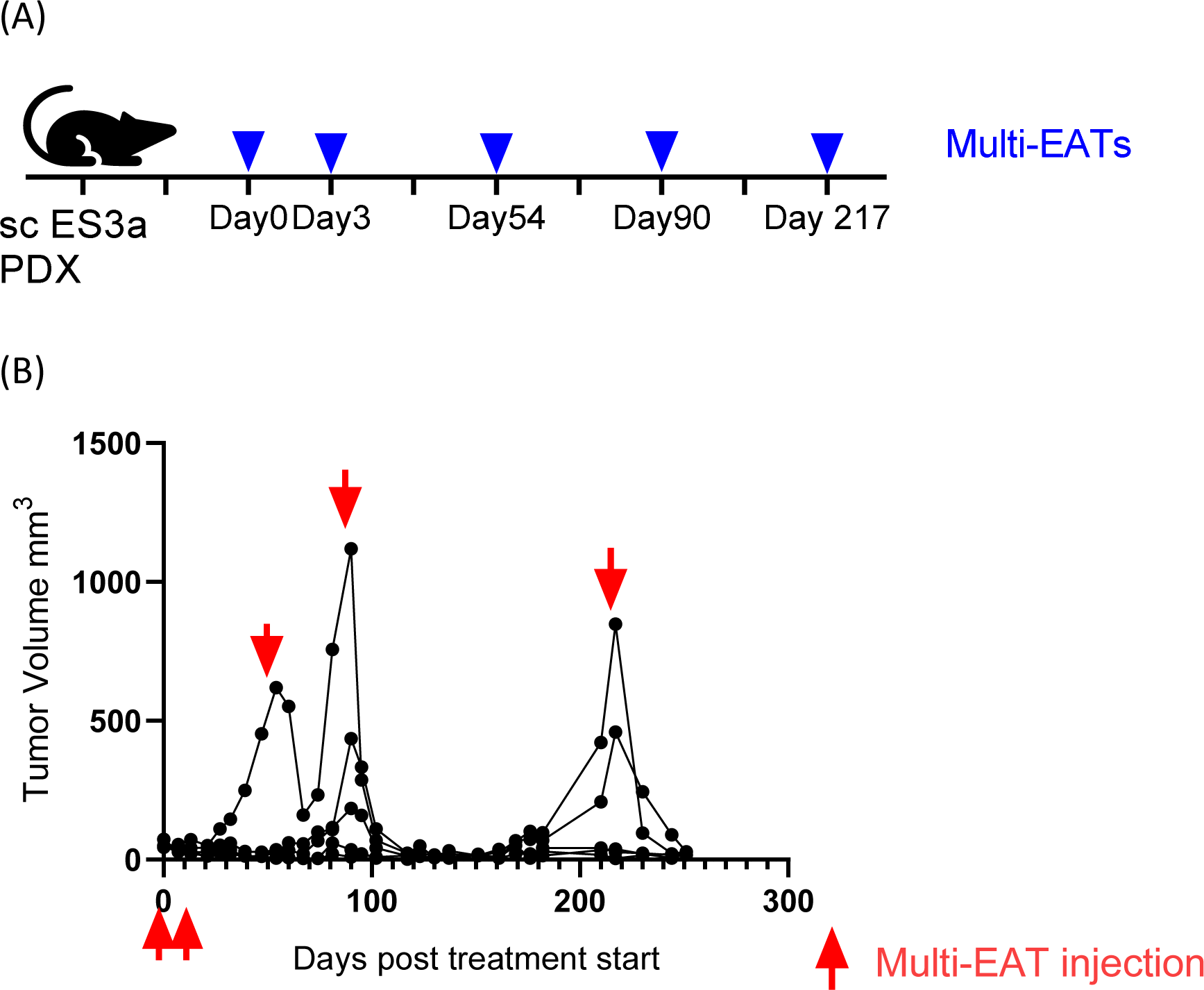
*In vivo* anti-tumor response of multi-EATs against relapsed tumors. (A) Overview of treatment. For multi-EAT therapy, 2x10^7^ of T cells were armed with 5 BsAbs (2µg of GD2-BsAb, 2µg of HER2-BsAb, 2µg of CD33-BsAb, 2µg of PSMA-BsAb, and 2µg of STEAP1-BsAb) and administered intravenously on day 0 and day 3 post-treatment. When tumors relapsed, identical doses of multi-EATs were given on day 54, day 90, and day 217, respectively. (B) *In vivo* anti-tumor response was monitored.

